# Alpha oscillations reflect similar mapping mechanisms for localizing touch on hands and tools

**DOI:** 10.1101/2022.09.01.506165

**Authors:** Cécile Fabio, Romeo Salemme, Alessandro Farnè, Luke E. Miller

## Abstract

Numerous studies have suggested that tools become incorporated into a representation of our body. A prominent hypothesis suggests that our brain re-uses body-based computations when we use tools. However, little is known about how this is implemented at the neural level. Here we used the ability to localize touch on both tools and body parts as a case study to fill this gap. Neural oscillations in the alpha (8-13 Hz) and beta (15-25 Hz) frequency bands are involved in mapping touch on the body in distinct reference frames. Alpha activity reflects the mapping of touch in external coordinates, whereas beta activity reflects the mapping of touch in skin-centered coordinates. Here, we aimed at pinpointing the role of these oscillations during tool-extended sensing. We recorded participants’ oscillatory activity while tactile stimuli were applied to either hands or the tips of hand-held rods. The posture of the hands/tool-tips was uncrossed or crossed at participants’ body midline in order for us to disentangle brain responses related to different coordinate systems. We found that alpha-band activity was modulated similarly across postures when localizing touch on hands and on tools, reflecting the position of touch in external space. Source reconstruction also indicated a similar network of cortical regions involved for tools and hands. Our findings strongly suggest that the brain uses similar oscillatory mechanisms for mapping touch on the body and tools, supporting the idea of neural processes being repurposed for tool-use.

**SIGNIFICANCE STATEMENT:** Tool use is one of the defining traits of humankind. Tools allow us to explore our environment and expand our sensorimotor abilities. A prominent hypothesis suggests that our brain re-uses body-based neural processing to swiftly adapt to the use of tools. However, little is known about how this is implemented at the neural level. In the present study we used the ability to map touch on both tools and body parts as a case study to fill this gap. We found that the brain uses similar oscillatory mechanisms for mapping touch on a hand-held tool and on the body. These results provide novel and compelling support to the idea that neural processes devoted to body-related information are re-purposed for tool-use.

## INTRODUCTION

Tools allow us to extend our physical body, therefore amplifying our sensorimotor abilities. It is theorized that tools become incorporated into a neural representation of our body (Head and Holmes, 1911; Maravita and Iriki, 2004; Martel et al., 2016) as tool use notably alters motor kinematics (Cardinali et al., 2009, 2016), representation of body metrics (Cardinali et al., 2011, 2016; Sposito et al., 2012; Miller et al., 2014), and representation of space around the upper limb (Berti and Frassinetti, 2000; Farnè and Làdavas, 2000). Alongside these studies, Miller et al. (2018) recently found that participants can accurately localize where an object touches the surface of a hand-held tool, thus using the tool as a sensory extension of their body. These behavioral effects prompted the hypothesis that the brain repurposes body-based neural processing to control and sense with a tool. However, evidence for this hypothesis is currently limited (Iriki et al., 1996; Miller et al., 2019; Fabio et al., 2022). Here we used tool-extended tactile localization as a case study to investigate the brain repurposing of body-based neural computations during tool use.

Comparing touch localization on hands and tools requires a firm grasp on the underlying neural computations that map touch on the body (Miller et al., 2022). At the level of neural oscillations, touch on the skin leads to a desynchronization of power in two main low-frequency bands: alpha (8-13 Hz) (Salmelin and Hari, 1994; Salenius et al., 1997; Cheyne et al., 2003; Neuper et al., 2006; Haegens et al., 2014) and beta (15-25 Hz) (Salmelin and Hari, 1994; Pfurtscheller et al., 2001; Neuper et al., 2006). These frequency bands have been implicated in tactile localization within two types of computational spatial codes (Heed et al., 2015): Beta activity reflects encoding in anatomical coordinates (Buchholz et al., 2013, 2011; Schubert et al., 2019, 2015), which correspond to the position of touch on the skin; Alpha activity reflects encoding in external coordinates (Buchholz et al., 2013, 2011; Ruzzoli and Soto-Faraco, 2014; Schubert et al., 2019, 2015), which correspond to the position of the touch in the egocentric space around the body.

Typically, the processes behind these spatial codes have been disambiguated by crossing the hands over the body midline: When crossed, the right hand (anatomical coordinates) is located in the left hemispace (external coordinates), thus creating left-right conflict that affects behavioral performance across several localization tasks (Yamamoto and Kitazawa, 2001; Shore et al., 2002; Heed and Azañón, 2014). This conflict is often emphasized by attention cueing paradigms, as mapping touch using external coordinates may require orienting spatial attention (Heed and Röder, 2010; Eardley and van Velzen, 2011; Schubert et al., 2019).

There is reason to believe that localizing touch on a tool may involve similar neurocomputational mechanisms. At a behavioral level, touch on hand-held tools can be localized extremely accurately (Miller et al., 2018) and is similarly impaired when the tips of the tools are crossed over the midline while the hands remain uncrossed (Yamamoto and Kitazawa, 2001; Yamamoto et al., 2005). Our previous EEG study showed the involvement of alpha activity in tool-extended tactile localization with sources in a network of parieto-frontal areas involved in tactile and spatial processing (Fabio et al., 2022). However, the nature of the reference frame(s) underlying the observed alpha activity remains unclear.

To fill this gap, here we investigated whether the oscillatory mechanisms involved in localizing touch on the body are repurposed when localizing touch on a tool. To determine this, we characterized and compared the reference frames reflected in alpha- and beta-band oscillatory activity, during body-based and tool-extended tactile localization. We used EEG to record oscillatory activity of participants performing a cued tactile localization task on their hands and on hand-held tools while manipulating their posture (crossed vs uncrossed). When tool-tips were crossed over the midline, the hands holding the tools were always uncrossed, allowing us to directly compare tool-based and body-based reference frames.

## MATERIALS AND METHODS

### Participants

20 right-handed participants (mean age: 24.8 years; range: 18-32 years, 10 males), free of any known sensory, perceptual, or motor disorders, volunteered to participate in the experiment. We chose this sample size in accordance to previous studies about somatosensory oscillations (Haegens et al., 2011; Schubert et al., 2015; Fabio et al., 2022). The experiment was performed in accordance with the ethical standards laid down in the Declaration of Helsinki (2013) and all participants provided written informed consent according to national guidelines of the ethics committee (CPP SUD EST IV RCN: 2010-A01180-39).

### Setup

The experiment was divided in two sessions wherein touch was delivered to either the participants’ hands (Hand condition) or to hand-held wooden rods (Tool condition); the setup was similar for both conditions. Throughout the experiment, participants sat in an adjustable chair in front of a table. In the Hand condition, they placed their forearm on the table, positioned either in an uncrossed or in a crossed posture (alternated blockwise, order counterbalanced across participants) with each index finger resting on a support at the edge of the table (Fig. 1A). In the Tool condition, participants’ arms were placed on adjustable armrests and they hold a 50cm long wooden rod in each hand, either in an uncrossed or in a crossed posture (alternated blockwise, order counterbalanced across participants). Notably, in the Tool condition, only the tool-tips -but not the hands-crossed theirbody midline (Fig. 1B)—the overall posture of the upper limb was relatively unchanged when tools were crossed or uncrossed. Depending on the condition, the index fingertips or the tip of each tool rested on a foamed stabilizing support at the edge of the table. When crossed, the rods were raised one above the other in order to avoid their contact during stimulation.

**Figure 1.**
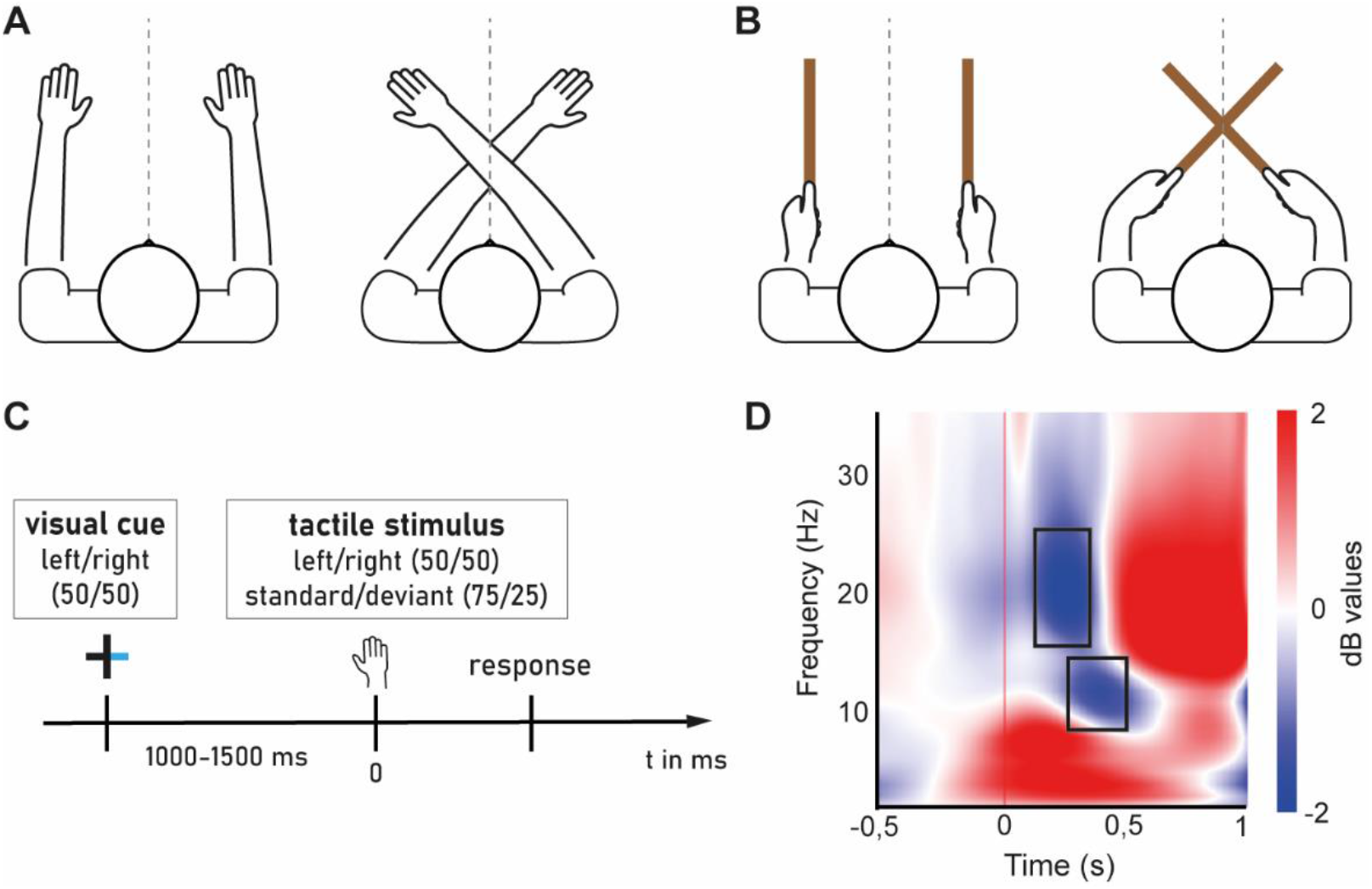
Experimental setup and paradigm. Participants (n=20) performed a tactile detectiontask for touches applied on two Surfaces: (A) when applied on Hands, participants hold their hands either in an Uncrossed posture (left) or a Crossed posture (right). (B) when applied on Tools, participants hold tools either in an Uncrossed posture (left) or a Crossed posture (right) where only the tool-tips crossed over the body midline (gray dotted line). (C) Trial structure of the tactile discrimination task. Tactile stimulation corresponds to the time zero. (D) Total oscillatory activity of the post-stimulation period over contralateral somatosensory cortex (electrode C3). Modulations are displayed as compared relative to baseline (−500 to −100 ms). Selected time windows for analysis are represented by grey rectangle for each frequency band: 250-500 ms for alpha and 150-350 ms for beta.

During the localization task, participants fixated on a central cross (2 cm wide) that was displayed on a 16’’ monitor in front of them and aligned with their body midline. Two solenoids (Mecalectro 8.19-.AB.83; 24 V, supplied with 36 W) were used to contact either index fingers or tool-tips, both with a disc surface of 4 cm to ensure uniform and consistent contact. Behavioral responses were made with a foot pedal (Leptron Footswitch 548561) that was placed underneath the left foot in half of the experiment, and under the right in the other half (alternated condition-wise, order counterbalanced across participants).

We took two approaches to mitigate the presence of auditory feedback from the solenoids. First, noise-cancelling earphones (Bose QuietComfort 20) playing white noise were used to mask the sound created by the solenoids. Further, to avoid any remaining auditory cue, each solenoid had another (decoy) solenoid placed on the opposite side, not in contact with the participants ‘hands or tools, that were both activated synchronously. All solenoids were mounted on adjustable tripods. Visual feedback was prevented by covering the table with a white cardboard.

### Experimental paradigm

Participants performed a tactile spatial discrimination task divided in two sessions depending on the surface stimulated (Hand or Tool). In this task, participants were cued to pay attention to one side of external space. They then had to detect a deviant stimulus (double tap) presented to a hand/tool in this cued side, while ignoring stimuli in the uncued side of space. Importantly, posture was manipulated to disentangle the involvement of different reference frame transformations. Each participant completed both sessions of the experiment on separate days (counterbalanced). Each session started with a practice block of 42 trials in both postures (Uncrossed and Crossed) to ensure they correctly understood and complied with task instructions.

At the beginning of each trial the central cross blinked to indicate the start of the trial. Then one side of the cross briefly turned blue (for 50 ms) to indicate which side of space (left or right, equal probability) participants had to attend to (Fig. 1C). After a variable delay (between 1000–1500 ms; randomly chosen from a uniform distribution), tactile stimulation was applied on participants’ right or left finger (Hand session) or on the tip of the right or left rod (Tool session). Note that this was independent of the cued side. Tactile stimuli were either frequent standard stimuli (solenoid raised once for 50 ms, including rise time and surface contact; probability of 0.75), or rare deviant stimuli (solenoid raised twice in a row for 50 ms separated by a 75 ms gap; probability of 0.25) presented with an equal probability in a random sequence to the left and the right. Participants had to respond as fast and accurately as possible using the foot pedal to rare tactile deviants presented to the cued side (“targets”, probability of 0.125), and to ignore standard stimuli at the attended side, as well as all stimuli presented to the other side. The experiment consisted of two sessions of 10 blocks, half Crossed, half Uncrossed. Each block included 60 standard trials and 20 deviant trials. The analysis included only trials in which standard stimuli were presented and in which, accordingly, no response was required, for a total of 600 trials per sessions. Participants complied with instructions, as evidenced by their high accuracy in each posture and for both surfaces (all>96%).

### EEG recording

EEG data were recorded continuously using a 65 channel ActiCap system (Brain Products). Horizontal and vertical electro-oculograms (EOGs) were recorded using electrodes placed below the left eye, and near the outer canthi of the right eye. Impedance of all electrodes was kept at <20 kΩ. FCz served as the online reference. EEG and EOG signals were low-pass filtered online at 0.1 Hz, sampled at 2500 Hz, and then saved to disk. Stimulus presentation and behavioral response collection were performed using MatLab on the experimental control computer, which was synchronized and communicated with the EEG data recording system. The trial events were sent to the EEG data recording system via a parallel port.

### Pre-processing of the EEG data

EEG signals were preprocessed using the EEGLab Toolbox (Delorme and Makeig, 2004). The preprocessing steps for each participant were as follows: for each session, participants’ five blocks in uncrossed posture, followed by their five blocks in crossed posture, were appended into a single dataset. The signal was resampled at 500 Hz and high-pass filtered at 0.1 Hz. Faulty channels were interpolated using a spherical spline. We then epoched data into a time window of 3.5 seconds, 1 second before and 2.5 seconds after the cue and baseline corrected using the period from −500 ms to −100 ms before the cue as baseline. Next, we removed signal artifacts with two steps: first, we removed eye blinks and horizontal eye movements from the signal using independent components analysis (ICA (Delorme and Makeig, 2004)) and a semi-automated algorithm called SASICA (Chaumon et al., 2015). We excluded every trial when participants made a response (whether it was a correct or an incorrect answer) or accidentally released the foot pedal, so that we only kept trials free of any motor activity. We re-epoched the data around the tactile stimulation (time zero), from −1.5 second before to 1 second after, leading to 2.5s epochs, and used the period from −500 ms to-100 ms before the hit for baseline correction. We then manually rejected trials that were contaminated by muscle artefacts or other forms of signal noise. In total, this led to a mean exclusion of 52.6 trials per participant (range: 2–205). Next, we used the EEGLab function *pop_reref* to add FCz (the online reference) back into the dataset and re-referenced the data to the average voltage across the scalp. Finally, for a better signal-to-noise ratio, we swapped the electrodes order of all the trials where the left hand or tool has been touched, so that all trials would now be in reference to the right hand or tool being touched (Buchholz et al., 2013; Schubert et al., 2019).

### Time-frequency decomposition

Time-frequency decomposition was performed using the open source toolbox Brainstorm (Tadel et al., 2011) in Matlab. The raw signal of each epoch was decomposed into frequencies between 1–35 Hz (linearly spaced) using Complex Morlet wavelets with a central frequency of 1 Hz and a full-width half maximum of 3 s. These parameters were chosen to ensure that our time-frequency decomposition had good spectral and temporal resolution within the chosen frequency range. Signal was then normalized in decibel (dB) to its ratio with the respective channel mean power during a baseline period ranging from −500 to −100 ms before tactile stimulation, which ensured that no post-stimulus activity contributed to the baseline normalization.

### EEG data analysis

Alpha- and beta-band activity were defined here as 8–13 Hz and 15–25 Hz respectively. Based on the observation of the oscillatory temporal dynamics during the time period after the hit (collapsed across all participants and conditions, see Figure 1D), we selected two time windows for analysis for each of our frequency band of interest: 150-350 ms for beta-band, and 250-500 ms for the alpha-band. This is a bias-free method for choosing time windows upon which running the analysis (Cohen, 2014).

Our experimental design had three factors that could be used in our analysis: Surface (Hand or Tool), Posture (Uncrossed or Crossed) and Attention (Unattended or Attended). Our initial analysis included all factors. The main effect of each factor was calculated by averaging the power of each frequency band, as well as the time points within the given time window before comparing the scalp topography of the relevant levels (e.g. Uncrossed vs. Crossed for main effect of Posture) using a cluster-based permutation test ((Maris and Oostenveld, 2007); two-tailed, cluster-level significance threshold of 0.05 and 1000 permutations run). Interaction effects were assessed via subtraction across conditions. For example, take the three-way interaction between Attention, Posture, and Surface. We first calculated the differences between unattended and attended stimulation for each posture separated by surface (e.g. Hand crossed unattended – Hand crossed attended). We then subtracted these differences for each surface before comparing them for each frequency band (average power) in their respective time windows (average time points; CBPT, same parameters). For all two-way interactions (e.g. Attention x Posture) we first collapsed across the unused condition (Surface); we then calculated the difference between levels of the first factor (Unattended – Attended) for each levels of the second factor (Uncrossed, Crossed). In a secondary analysis, we analyzed both surfaces separately.

### Source reconstruction

We followed up significant interactions with source reconstruction for each epoch to estimate which brain regions were involved. This was done using the open source toolbox Brainstorm (Tadel et al., 2011). First, a head model was computed using OpenMEEG BEM model (Gramfort et al., 2010). A noise covariance matrix for every participant was computed over a baseline time window of −500 to −100 ms before stimulation. Sources were then estimated using the Standardized low resolution brain electromagnetic tomography (sLORETA(Pascual-Marqui, 2002)) approach with unconstrained dipole orientations across the surface. We then performed time-frequency decomposition on the source files to localize significant power modulations in the alpha-band. The signal at each vertex was decomposed into the mean of frequencies going from 8 to 13 Hz using Complex Morlet wavelets with a central frequency of 1 Hz and a full-width half maximum of 3 s. The signal was then normalized in decibel (dB) to its ratio with the respective channel mean power during a baseline period ranging from −500 to −100 ms before tactile stimulation.

The interaction between Attention and Posture was assessed in a similar manner as previously described. We calculated the difference in alpha power between unattended and attended stimulation for each posture and compared them in the chosen time windows (average time points) using Cluster-based permutation ((Maris and Oostenveld, 2007); two-tailed, cluster-level significance threshold of 0.01 and 1000 permutations run).

## RESULTS

### Similar oscillatory correlates for tactile localization on hands and tools

The present study investigated whether there were different oscillatory correlates for localizing touch on hands and on tools, with a specific focus on the external remapping of touch. Therefore, the contrasts containing the factor Surface (Hand, Tool) are of particular interest for us. This includes one three-way interaction (Attention x Posture x Surface), two two-way interactions (Attention x Surface, Posture x Surface) and a main effect of Surface. We therefore first determined whether there were statistically significant differences in the oscillatory power of alpha and beta for these contrasts. Significant differences in these contrasts would indicate that tactile-localization modulated oscillatory power differently between the hand and the tool.

Using a cluster-based permutation test (CBPT, α-range = 0.05), we found no significant interactions or main effects with the factor Surface in either the alpha or beta-band (Table 1, CBPT: p>0.05).

**Table 1.** Contrasts performed on scalp topographies of mean oscillatory power; CBPT, p-value of all clusters found. ‘/’ indicates no cluster with a p-value below 1.

| Contrasts | Frequency band | p-value |
| --- | --- | --- |
| Posture | Alpha | 0.316 |
|  | Beta | / |
| Attention | Alpha | 0.012 |
|  | Beta | 0.046 |
| Surface | Alpha | 0.276 & 0.346 |
|  | Beta | 0.164 & 0.359 |
| Posture x Surface | Alpha | / |
|  | Beta | / |
| Attention x Surface | Alpha | / |
|  | Beta | 0.342 |
| Attention x Posture | Alpha | 0.006 & 0.049 |
|  | Beta | 0.194 |
| Attention x Posture x Surface | Alpha | 0.242 |
|  | Beta | / |

### Spatial attention modulates tactile processing according to posture

Given the lack of any statistical differences between oscillations while localizing touch on hands and tools, we next investigated the surface-independent effects of posture on oscillations. A significant effect containing the factor Posture (uncrossed vs. crossed) would suggest processing related to an external reference frame. We did not observe a general main effect of Posture for alpha or beta power (Table 1, CBPT: p>0.05).

We did, however, observe a significant interaction effect between Attention and Posture for power in the alpha band in both hemispheres (left: p-value=0.049 &right: p-value=0.006). The two significant clusters were localized above parieto-occipital channels in their respective hemisphere (Fig. 2A). As can be seen in Figure 2A, touch led to widespread alpha desynchronization across centro-posterior channels, which was increased when attention was directed to the touched side (Main effect of attention: p<0.05, see Table 1). Crucially, we observed that crossing the surface (hand or tool) shifted the topographic distribution of alpha desynchronization, but did so differently for the attended and unattended conditions (lower panels of Figure 2A).

**Figure 2.**
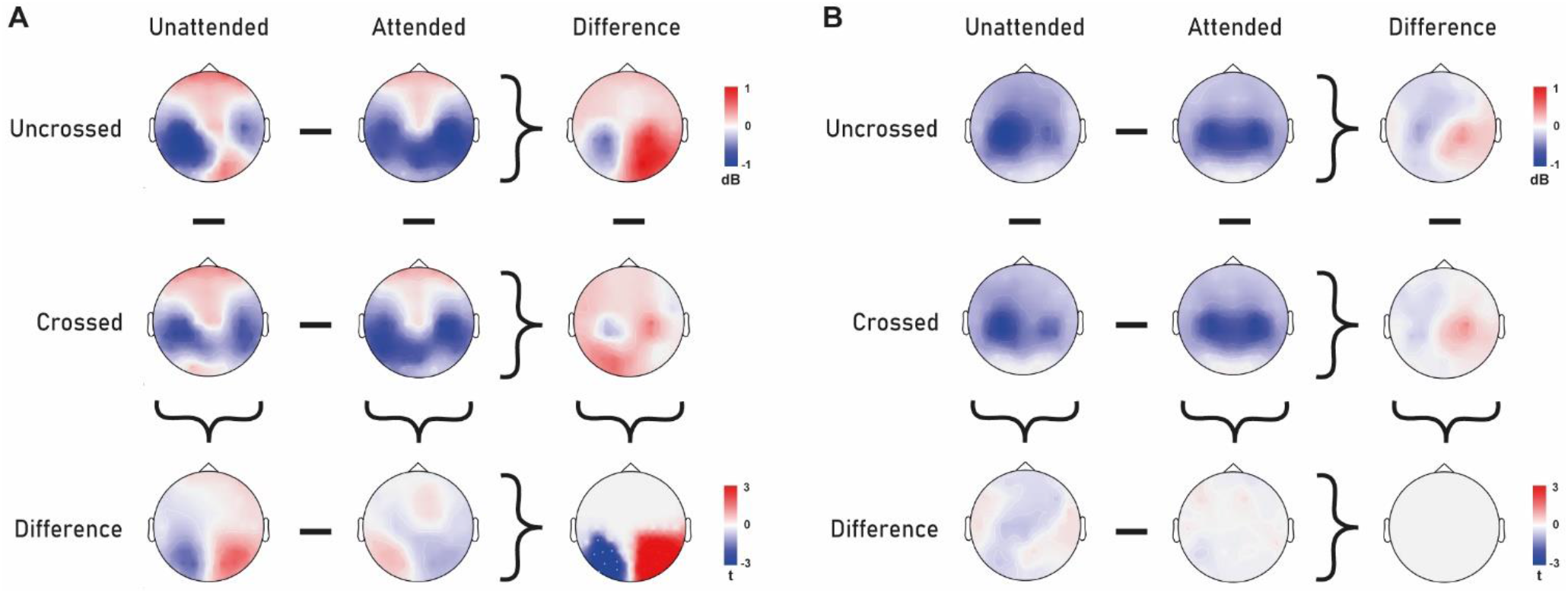
Alpha and beta activity after tactile stimulation. (A) Topographies of alpha-band activity (8–13 Hz, 250 to 500 ms) in uncrossed (1^st^ row) and crossed (2^nd^ row) posture following unattended (1^st^ column) and attended stimuli (2^nd^ column). Difference topographies for attention effects in uncrossed and crossed posture (3^rd^ column), and for posture effects following attended and unattended stimuli (3^rd^ row). Bottom-right corner: topography of the interaction between attention and posture. (B) Topographies of beta-band activity (15-25Hz, 150 to 300ms) in uncrossed (1^st^ row) and crossed (2^nd^ row) posture following unattended (1^st^ column)and attended stimuli (2^nd^ column). Difference topographies for attention effects in uncrossed and crossed posture (3^rd^ column), and for posture effects following attended and unattended stimuli (3^rd^ row). Bottom-right corner: topography of the interaction between attention and posture. Data are displayed as if stimuli always occurred on the anatomically right hand or the tool held in the right hand, so that the left hemisphere is contralateral to tactile stimulation in a skin-based reference frame, independent of posture.

In the beta band, we observed a bilateral decrease of power over central channels that was independent of Posture. We found that the desynchronization of beta was more bilateral in the Attended condition (Main effect of attention: p<0.05, see Table 1). However, we did not find a significant interaction between Attention and Posture in the beta band (Fig. 2B) and thus do not consider this frequency band in later analyses.

### Similar topography of alpha desynchronization is observed for hands and tools

We then analyzed each Surface separately, in order to further explore the localization processes for touch on hands and on tools. Visual inspection of the scalp distribution of alpha activity for each Surface reveals a striking resemblance between Hand and Tool (Fig. 3A&B). For each condition, alpha power modulation is almost identical between the two surfaces, with patterns reflecting what we observed when Surfaces were collapsed (Fig. 2A). This underscores the inference that the neural processes underlying localizing touch on each surface are similar.

**Figure 3.**
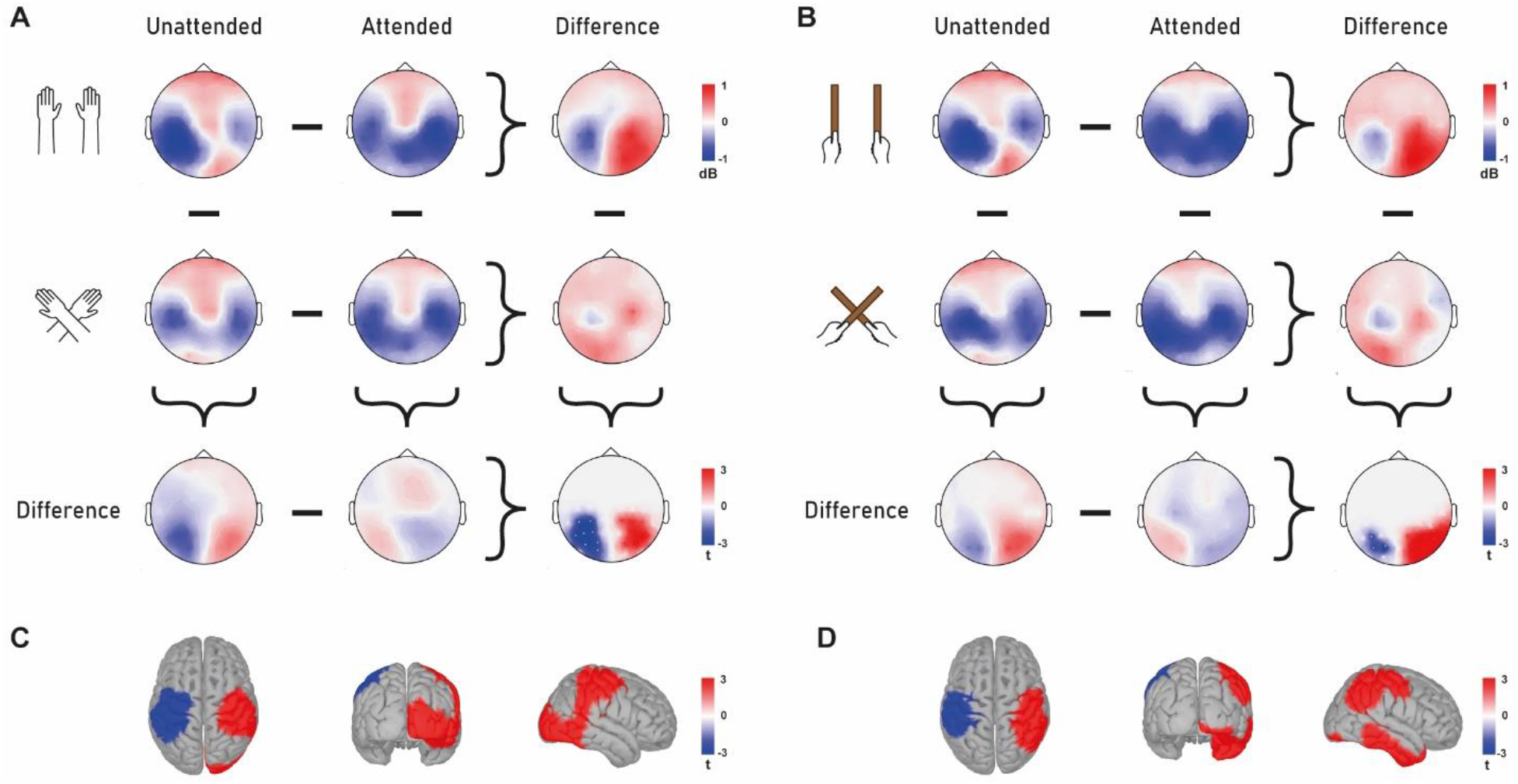
Alpha activity following tactile stimulation on the hand and on the tool. (A) Topographies of alpha-band activity (8–13 Hz, 250 to 500 ms) when tactile stimuli happened on the hand, with uncrossed (1^st^ row) and crossed (2^nd^ row) hands following attended (1^st^ column) and unattended (2^nd^ column) stimuli. Difference topographies for attention effects with uncrossed and crossed hands (3^rd^ column), and for posture effects following attended and unattended stimuli (3^rd^ row). Bottom-right corner: topography of the interaction between attention and posture. (B) Topographies of alpha-band activity (15-25Hz, 150 to 300ms) when tactile stimuli happened on the tool, with uncrossed (1^st^ row) and crossed hands (2^nd^ row) following attended (1^st^ column) and unattended (2^nd^ column) stimuli. Difference topographies for attention effects with uncrossed and crossed tools (3^rd^ column), and for posture effects following attended and unattended stimuli (3^rd^ row). Bottom-right corner: topography of the interaction between attention and posture. (C) Source reconstruction of the interaction effect between attention and posture for tactile stimulation on the hand. (D) Source reconstruction of the interaction effect between attention and posture for tactile stimulation on the tool. Data are displayed as if stimuli always occurred on the anatomically right hand or tool held in the right hand, so that the left hemisphere is contralateral to tactile stimulation in a skin-based reference frame, independent of posture.

We then calculated the interaction between Attention and Posture for each surface separately. We found a similar pattern of interaction between hand and tool: For stimulation on the hand, we obtained two clusters, one in each hemisphere (CBPT, left: p= 0.036 & right: p=0.074). We also obtained two clusters with similar distribution when stimulation happened on the tool (CBPT, left: p=0.195 & right: p=0.022). While not all clusters reached statistical significance for each surface, their overall distribution corresponded well to the interaction between Attention and Posture observed when surfaces were collapsed (Fig. 2A) and therefore displaying a comparable pattern of reference-framed based oscillatory processing.

We next identified the cortical sources underlying this interaction effect for each surface. We observed similar sources for localization on hands and tools: The interaction effect was significant throughout sensorimotor regions, including the primary somatosensory (SI), primary motor cortices (MI) and posterior parietal regions contralateral to the stimulated hand (Fig. 3C, p = 0.046) and tool (Fig. 3D, p = 0.024). In the hemisphere ipsilateral to the stimulated hand, cortical sources included the same sensorimotor frontoparietal regions as well as the occipitotemporal cortex (Fig. 3C, p = 0.006). The interaction effect in the hemisphere ipsilateral to the stimulated tool also spread over the same frontoparietal regions (Fig. 3D, p = 0.006) but, in contrast with the hand, also included a larger portion of the temporal cortices (Fig. 3D, p = 0.019). In general, nearly identical sources were found for mapping touch on either hands or tools.

## DISCUSSION

The present study was designed to identify the spatial codes used for localizing tactile stimuli delivered on hand-held tools and to compare them with those typically used when localizing tactile stimuli applied on hands. To this end, we used EEG in a cued tactile localization task whereby we manipulated hand and tool posture (crossed vs. uncrossed). We found a remarkable similarity of alpha and beta power modulation following touch between the two surfaces. Importantly, there was no main effect of posture for either surface, but a significant interaction between attention and posture that was selective for the alpha-band. This effect was also similarly distributed across channels for hand and tool. Furthermore, source localization of this effect for both surfaces revealed that comparable cortical networks were involved. Overall, these findings provide evidence that similar neurocomputational mechanisms are used by the brain to process touch location on the hand and on a hand-held tool. These mechanisms are reflected by alpha activity when manipulating posture, suggesting the use of external coordinates.

### Touch on hands and tools rely on shared oscillatory mapping mechanisms

The main result of this study is that localizing touch on hands and tools involves similar oscillatory correlates. Indeed, not only no significant difference was found between oscillatory power of alpha and beta-band between surfaces (see Table 1), but most notably their scalp topographies were almost identical between hand and tool when observed separately (Fig. 3A&B for alpha, beta not shown). These results appear to be consistent with the centuries-old proposal of tool embodiment (Head and Holmes, 1911). Incorporation of a hand-held tool into body representation may indeed consist in repurposing the neural mechanisms that process body-related sensory information for processing information originating from the tool. Until now, neuroscientific evidence for this proposition has been scarce, since the majority of evidence comes from behavioral studies and from paradigms that only measure the effects that tool-use induced on subsequent perceptual or motor measures (Berti and Frassinetti, 2000; Farnè and Làdavas, 2000; Cardinali et al., 2009, 2011, 2016; Sposito et al., 2012; Miller et al., 2014; Forsberg et al., 2019). For example, initial evidence of online repurposing comes from Iriki and colleagues’ work who measured from macaque monkeys’ multisensory postcentral neurons during tool-use and observed an expansion of the visual portion of their receptive field to encompass the tool (Iriki et al., 1996). At the behavioral level, online remapping of space was observed during tool-use by Berti & Frassinetti (2000), but their neuropsychological approach could not provide indications as to which mechanisms are at play. The present study overcame this limitation by measuring oscillatory activity underlying reference frame transformations for touch on hands and tools.

Our findings are consistent with previous studies on the neural correlates of tool sensing. We previously recorded EEG activity of human participants during a tactile localization task on a hand-held tool, therefore directly observing online tool-use. We identified a modulation of alpha power dependent of contact location (Fabio et al., 2022), suggesting that it is a signature of tool-extended tactile localization. Consistent with these previous results, here we also found that posture modulated alpha activity dependently of attention: for both surfaces, interaction effects were localized in two parieto-occipital clusters, one in each hemisphere (Fig. 3A&B). We found some differences in the distribution of interaction effect of Attention and Posture between Surfaces. The right occipital cortex was notably activated for the hand, whereas the inferior temporal cortex was activated for the tool. This could be explained by the effect of attention on actively shaping and enhancing spatial representations in the ventral visual pathway (Kay et al., 2015). Besides this difference, the source reconstruction of the alpha modulation was largely comparable between surfaces (Fig. 3 C&D). This new evidence adds to our previous ERP study (Miller et al., 2019) on tool-extended sensing: touches on the tool and on the arm led to similar stages of cortical processing as well as similar sources involved.

In sum, the remarkable similarity that we found for oscillatory processes for tactile localization on the hand and on the tool suggests that in order to localize a contact happening on a hand-held tool, the human brain repurposes neural mechanisms dedicated to body-related processes to perform the same function with a tool.

### Alpha rhythm reflects external spatial coding for touch on hands and tools

Crossing limbs is a well-established method to tease apart localizing processes in external and skin-based coordinates. In this respect, previous electrophysiological studies have linked alpha oscillations to a use of external coordinates and beta to skin-based coordinates (Buchholz et al., 2013, 2011; Ruzzoli and Soto-Faraco, 2014; Schubert et al., 2019). Here, we manipulated hands and tools posture to characterize and compare the cortical oscillations reflecting the crossing effects emerging from hands and tools. This is especially of note for the tool, since only the tool-tips crossed the body midline while the hands stayed in their respective hemispace (see Fig. 1A). Therefore, any effect observed for crossing when touch is on the tool surface would reflect the remapping of touch on the tool, not the hands.

Consistent with previous findings, we observed that modulation of alpha activity following posture was dependant on attention, which wasn’t the case for beta, supporting the involvement of the alpha band in external processing. Crucially, this modulation of the alpha activity was independent of whether touch localization processes concerned the hands or the tools, as exemplified in the near-identical scalp topographies and significant posterior clusters for both surfaces (Fig. 3A&B). Our previous study (Fabio et al., 2022) also found an involvement of alpha oscillations in the encoding of touch location on tools. While this suggested the encoding of an external spatial code, the paradigm we used did not manipulate posture and was therefore equivocal in these regards. However, the present results indeed support the proposition made in Fabio et al. (2022) that touch on a tool is primarily coded in an external reference frame.

Furthermore, encoding touch localization in external coordinates on the hand and on the tool involves a similar cortical network. Source reconstruction of the alpha coding of external space identified several regions throughout the parietal and frontal cortices in both hemispheres. This included primary somatosensory and motor cortex. Importantly, the posterior parietal sources, also shared by the two surfaces, have previously been implicated in the processing of touch in external space (Bremmer et al., 2001; Lloyd et al., 2003; Avillac et al., 2005; Azañón et al., 2010). Parietal alpha oscillatory activity indeed appears to play a crucial role in this process (Buchholz et al., 2011; Ruzzoli and Soto-Faraco, 2014; Schubert et al., 2019). To summarize, we found that alpha band indexes the spatial coding of touch in an external reference frame, this effect being independent of whether touch was localized on the hand or a hand-held tool.

### Alpha-based coding of external coordinates may depend on attention

Unsurprisingly, we found attention to modulate the overall oscillatory activity of both alpha and beta bands (see Table 1; (Sauseng et al., 2005; van Ede et al., 2010, 2011)). However, it is noteworthy that posture manipulation itself wasn’t sufficient to affect oscillatory activities; the modulation of alpha power following the crossing of the hands/tools was dependant on attention. This finding suggests that external spatial of touch was dependent of certain attentional processes. Alpha oscillations have indeed been implicated in tactile spatial attention (Bauer, 2006; van Ede et al., 2011, 2014; Bauer et al., 2012). Post-touch alpha oscillations may thus reflect the orienting of attention in external space (Ossandón et al., 2020).

This involvement of attentional processes in spatial coding was reflected in the scalp topographies and source localization of our interaction effect, which was mostly localized in somatosensory (Mima et al., 1998; Bauer et al., 2012) as well as posterior regions of the cortex, for the hand as well as the tool (Fig.3). This pattern of results is fitting with previous studies about the modulatory effects of attention on tactile ERP (García-Larrea et al., 1995; Eimer and Forster, 2003). In particular, Eimer et al. (2003) suggested that different spatial coordinate systems may be used by separable attentional control processes with a posterior process operating on the basis of external spatial coordinates, whereas an anterior process is based primarily on anatomically defined spatial codes (Eimer et al., 2003). Since crossing the hands (and hand-held tools) mainly modulate the external coordinates of tactile processing, our tactile spatial localization task likely involved spatial attentional processes taking places in external coordinates. Along these lines, using a similar experimental paradigm, Yue et al. (2009) also found that ERPs to tactile stimuli presented at the tips of tools were modulated by spatial attention (Yue et al., 2009).

To conclude, we found that the brain uses similar oscillatory mechanisms for mapping touch on a hand-held tool and on the body. These results are in line with previous work and support the idea of that neural processes devoted to body-related information are being re-used for tool-use. Furthermore, alpha-band modulation followed the position of touch into external space. This is thus the first neural evidence that tactile localization on a hand-held tool involves the use of external spatial coordinates.

## Acknowledgments

This work was supported by the Agence Nationale de la Recherche [ANR-16-CE28-0015] Developmental Tool Mastery to A.F. & [ANR-19-CE37-0005] BLIND_TOUCH to A.F. and L.E.M; Fondation pour la Recherche Medicale Post-doctoral Fellowship SPF20160936329 to L.E.M. and by a Foundation Berthe Fouassier Doctoral fellowship to C.F. under the umbrella of the Fondation de France. All work was performed within the frame work of LABEX CORTEX [ANR-11-LABX-0042] of Université de Lyon. We thank Frederic Volland for his help constructing the experimental setup.

